# Establishing genetic tractability in *Jeongeupia sacculi* HS-3: disruption of *rpoS* hinders biofilm formation

**DOI:** 10.64898/2026.10.09.758012

**Authors:** Zachary Beddingfield, Isabel Calimlim, Christopher Zhang, Brian K Hammer

## Abstract

*Jeongeupia sacculi* (HS-3) is a novel *β*-proteobacterium with a unique multicellular life cycle. The genetic tractability and mechanisms underlying differentiation are unknown. Here, we adapt a method of homologous recombination for genetic modification in HS-3 and generate a disruption mutant of *rpoS*, which encodes the alternative sigma factor *σ*^S^ commonly associated with general stress response or major behavioral switches. We then characterize phenotypes associated with *rpoS* disruption. Under low-nutrient conditions, *rpoS* disruption reduced Congo red binding, consistent with *rpoS* acting as a regulator of low-nutrient response by upregulating production of biofilm-associated materials. Surprisingly, *rpoS* disruption did not significantly alter survival in response to other stressors in general stress response assays tested. Our work establishes a method for genetic manipulation in HS-3 and enables future investigation into the life cycle of HS-3, including whether *rpoS* has a role in differentiation triggered by low-nutrient response.

## Introduction

*Jeongeupia sacculi* (HS-3) is a *β*-proteobacterium isolated from a limestone cave wall on the Hirao karst plateau in Japan that exhibits a unique multicellular phase in its life cycle (1). When grown on a medium mimicking a nutrient-poor cave environment, an HS-3 colony forms as a filamentous biofilm of physically attached cells. A population of motile coccoid cells then arises within the colony, expanding and releasing into the water column upon submersion in water. While HS-3 was isolated and characterized by Mizuno et al.(1), little is known about its genetic basis. Consequently, the molecular mechanisms driving cellular differentiation, attachment, and dispersal remain unresolved. While better characterization of the genetics driving HS-3’s life cycle would contribute to our understanding of how emergent multicellular properties arise in prokaryotes, such work requires establishing HS-3 as a genetically accessible model. To enable these investigations, we developed effective cloning tools and successfully generated the first targeted mutant strain in this novel species. Although gene editing strategies have been well optimized in model organisms, genetic tractability varies substantially across species (2, 3). In non-model bacteria, genetic manipulation remains limited by organism-specific barriers to DNA uptake, defense mechanisms against foreign DNA, and host-dependent performance of selectable markers and reporter genes (4, 5). For example, plasmid persistence and plasmid-borne gene expression can vary among bacterial hosts (6, 7), and even strains carrying genes for natural competence may require optimization of transformation conditions for successful genetic modification (8). While Mizuno et al. demonstrated conjugation of an episomal, GFP-containing vector in HS-3, HS-3’s broader genetic tractability is unknown. To enable future investigation of the genetic mechanisms that regulate HS-3’s novel lifestyle, we set out to adapt existing genetic tools for targeted mutagenesis in HS-3.

Our approach built upon foundational cloning techniques, allowing us to successfully establish a streamlined protocol for targeted mutagenesis in HS-3. We chose to target HS-3’s *rpoS* gene, which encodes the alternative sigma factor *σ*^S^, for disruption by homologous recombination. While *σ*^S^ was originally understood to be a canonical master regulator of general stress response in *γ*-proteobacteria (9), further characterization has revealed it is also a regulator of environmentally induced physiological and lifestyle transitions (9, 10). In *Escherichia coli, σ*^S^ governs survival under oxidative, osmotic, and acid stress (11). In *Pseudomonas aeruginosa, σ*^S^ similarly governs oxidative and osmotic stress (12), but also quorum sensing (13), expression of virulence factors (12), and transition from a planktonic to a biofilm-associated lifestyle (14, 15). In *Vibrio cholerae, σ*^S^ activates mucosal escape response, triggering changes in gene expression causing *V. cholerae* to detach from and exit the host (16). Because *σ*^S^ is well-known to control a broad range of phenotypes across organisms, it provides an excellent entry point for gaining insight into the regulatory architecture of novel species. Further, given the consistent role of *σ*^S^ in triggering large changes to gene expression in response to stress and its association with various novel behavioral shifts, we reasoned *σ*^S^ likely plays a role in the complex life cycle of this environmental bacterium.

Here, we describe the successful creation of the first mutant of HS-3, constructed using homologous recombination to disrupt the *rpoS* gene. Throughout, the HS-3 disruption mutant *rpoS*::*kan*^R^ is designated *rpoS*^*−*^, and the strain designated WT here represents a variant of the parental wild-type strain that was created by random insertion of the identical *kanR* cassette into a chromosomal region that did not alter the phenotypes tested. The unmodified isolate is referred to as parental HS-3. We compare the survival of the WT and *rpoS*^*−*^ strains following exposure to several canonical stressors: hydrogen peroxide, ethanol, and elevated heat. We also measure each strain’s capacity to bind Congo red stain as a proxy for biofilm formation.

Our work demonstrates the genetic tractability of HS-3, providing an essential foundation for further investigation of the mechanisms of its multicellular differentiation. Further, we characterize the role of *σ*^S^ in HS-3, giving insight into the ways that *rpoS* can be co-opted for adaptation.

## Results

### *rpoS*^*−*^ is the first mutant of HS-3

Sequence analysis of the genome of HS-3 identified five genes encoding sigma factors, with the three-letter *rpo* mnemonic: *rpoD* (*σ*^70^), *rpoE* (*σ*^24^), *rpoH* (*σ*^32^), *rpoN* (*σ*^54^), and *rpoS* (*σ*^38^) (Fig. 1A). Since these sequences contain similar domains, we confirmed that the gene annotated as *rpoS* was a true homolog encoding *σ*^S^ by comparing the gene neighborhood and predicted protein domain architecture with those of reference RpoS homologs (Fig. 1B,C).

**Fig. 1.**
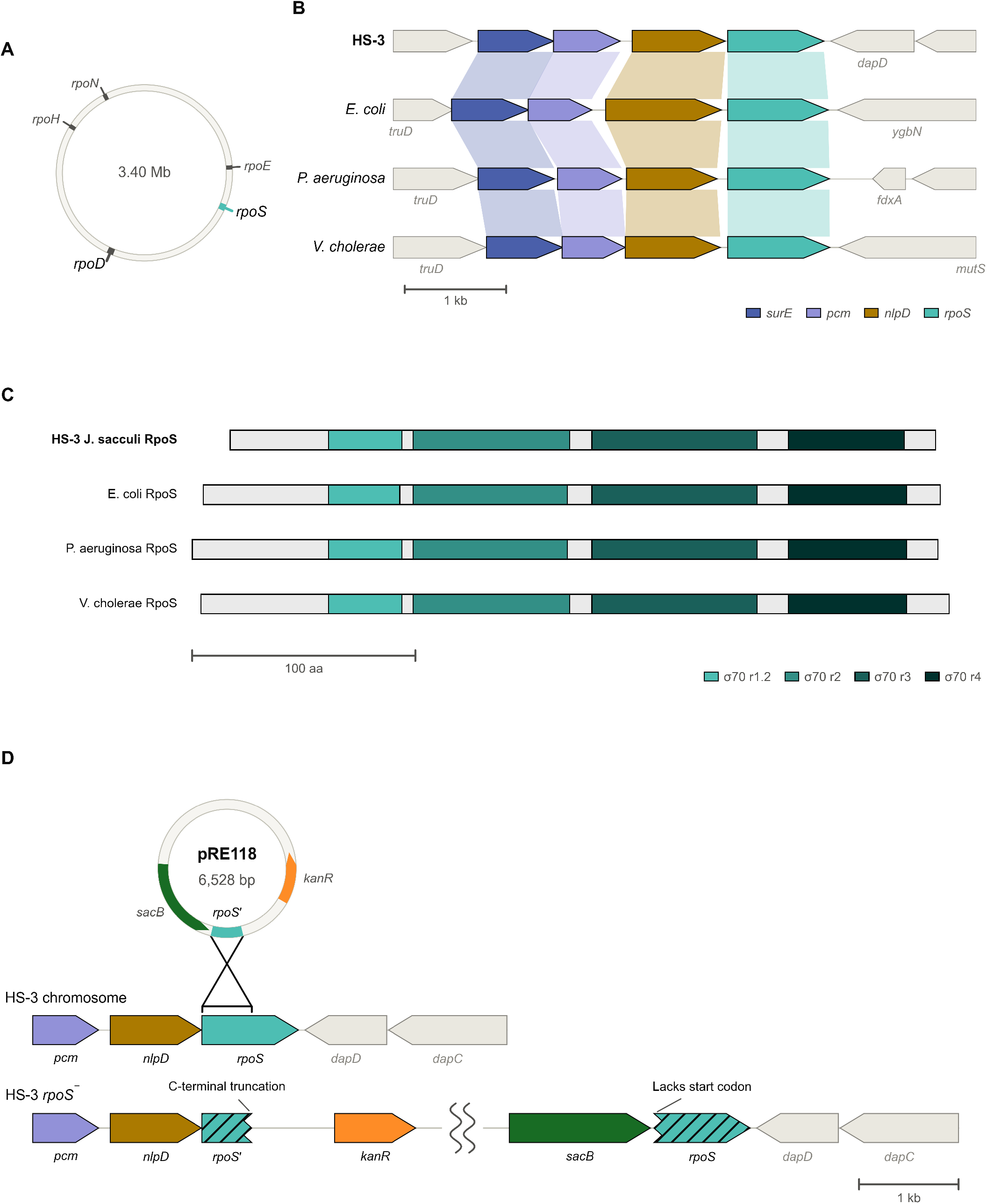
Characterization of HS-3 *rpoS* and cloning approach. (A) Chromosomal map of HS-3 depicting loci of sigma-factor genes. (B) Comparison of the genomic neighborhood surrounding HS-3 *rpoS* with those of three reference homologs, *Escherichia coli, Pseudomonas aeruginosa*, and *Vibrio cholerae*, reveals a conserved genomic neighborhood consistent with its annotation as *rpoS*. (C) Domain architecture of HS-3 RpoS compared with three reference homologs shows that HS-3 RpoS contains domains homologous to those in the reference proteins. (D) Depiction of mechanism of homologous recombination between *rpoS* and an internal fragment of *rpoS* lacking its native start codon encoded in a pRE118 plasmid to produce an *rpoS*::*kan*^R^ insertion mutant, referred to as *rpoS*^*−*^ in this study. The resulting mutant has both a truncated *rpoS* fragment and a copy lacking its native promoter and start codon.

At the domain level, *rpoS* is canonically comprised of conserved regions 1.2, 2, 3, and 4, whereas *rpoD* is comprised of these same regions and an additional conserved region 1.1 (17). Annotation by InterProScan and the Pfam 38.2 database confirmed that HS-3 *rpoS* carries the region 1.2, 2, 3, and 4 domains characteristic of *rpoS* while lacking the 1.1 domain that is characteristic of *rpoD* (Fig. 1C). Multiple sequence alignment plots of each domain are provided as Supplementary Figs. 1–4. Neighborhood and domain synteny of *rpoD, rpoH, rpoN*, and *rpoE* for HS-3 were also consistent with reference groups (Supplementary Fig. 5). Thus, the gene annotated as *rpoS* in HS-3 is indeed *rpoS* and not another sigma factor.

We chose homologous recombination with a suicide vector as our strategy for disrupting *rpoS*, given the unknown genetic tractability of HS-3 and well-established nature of the protocol. To identify resistance markers that could be used to select for suicide plasmid integration into HS-3, we first measured minimum inhibitory concentrations (MICs) for parental HS-3 grown in the presence of the antibiotics gentamicin, kanamycin, and chloramphenicol. HS-3 is susceptible to gentamicin (160 *µ*g/mL, *n* = 3), kanamycin (100 *µ*g/mL, *n* = 4), and chloramphenicol (10 *µ*g/mL, *n* = 4) (Supplementary Table 1). We chose kanamycin as a selectable marker for construction of the *rpoS*^*−*^ mutant. Previously, we attempted a strategy of deletion by homologous recombination for a different gene utilizing *sacB* counter-selection. However, we encountered difficulty isolating deletion mutants from those acquiring mutations in the *sacB* gene. To create an *rpoS* mutant without the requirement of counter-selection, we created a new plasmid construct for inactivation of *rpoS* by disruption. This construct carried a single 500 nt PCR-amplified internal fragment of *rpoS*. We successfully disrupted the *rpoS* gene by conjugating the suicide plasmid into HS-3 and then selecting for a single crossover event by plating on medium containing kanamycin. This produced a mutant in which the entire plasmid inserted within the *rpoS* gene (Fig. 1), as was confirmed by whole genome sequencing. This *rpoS*^*−*^ strain was compared with a control WT::*kanR* mutant, designated WT, for canonical RpoS-dependent phenotypes.

### *rpoS* disruption reduces Congo red binding in HS-3 grown on R2A

We speculated that attachment of HS-3 cells to one another and/or a cave wall might require formation of biofilms, which in many bacteria are under *σ*^S^ control (14, 17–19). Congo red (CR) staining is a common method used to assess biofilm-associated matrix production, including in *P. aeruginosa*, as CR binds exopolysaccharides that many bacteria use to attach to surfaces (20–22).

We plated single colonies of *rpoS*^*−*^ and WT HS-3 on both nutrient-rich Miller LB and R2A, a low-nutrient medium more likely to mimic stressful cave conditions (23–25), to assess the correlation of *rpoS* disruption and CR staining across different conditions. *Pseudomonas aeruginosa* PA14, which forms biofilms that are under control of RpoS, served as a control (14).

To compare Congo red staining quantitatively, we calculated a chromatic staining index, OD_chrom_, from colony images (Fig. 2B). OD_chrom_ is the difference between apparent optical densities in the green and red channels, each normalized by the surrounding agar. Since OD_chrom_ index increases with reflected light in the red channel and with absorption in the green channel, it works well for quantifying CR staining, which reflects red and absorbs green. Subtracting the channel-specific optical densities cancels multiplicative brightness changes that affect both channels equally, reducing sensitivity to spectrally neutral brightness variation and differences in biomass.

**Fig. 2.**
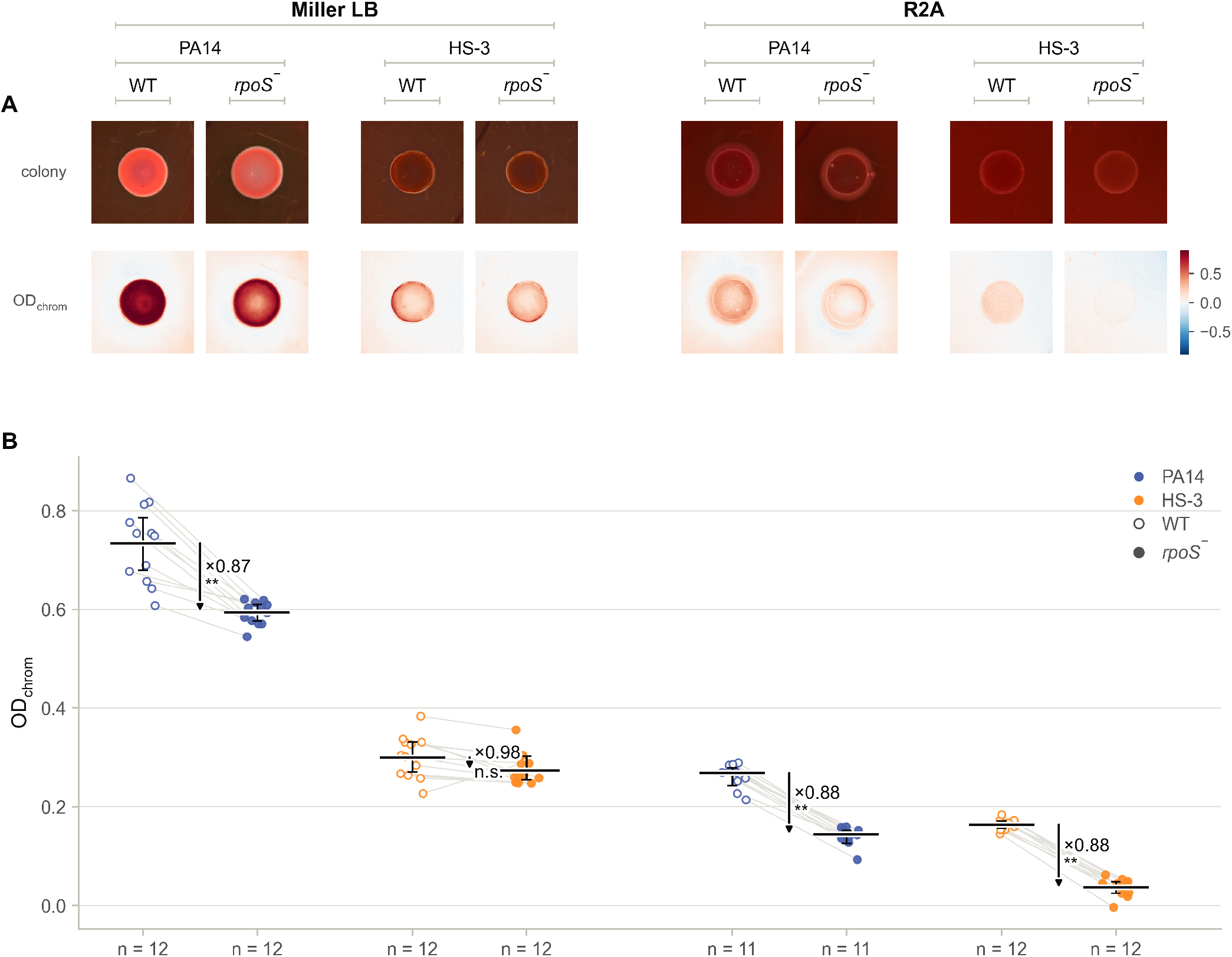
Effect of *rpoS* disruption on Congo red (CR) binding across media. (A) Representative RGB images and corresponding spatial OD_chrom_ maps from WT and *rpoS*^*−*^ strains of *Pseudomonas aeruginosa* PA14 and *Jeongeupia sacculi* HS-3 in Miller LB or R2A. (B) Measured OD_chrom_ values for WT (hollow) and *rpoS*^*−*^ (filled) colonies of PA14 (blue) and HS-3 (orange) by medium at 72 h. Disruption of *rpoS* significantly reduced CR staining in PA14 on both media. In HS-3, a significant reduction was detected on low-nutrient R2A but not on Miller LB. Points represent individual colonies, and lines connect colonies from the same petri dish. Horizontal bars mark Hodges-Lehmann location estimates for each group, and error bars show their unadjusted two-sided 95% Wilcoxon signed-rank CIs. Each arrow is anchored at the WT location estimate, and its vertical displacement represents the paired Hodges-Lehmann estimate of the within-dish difference, Δ = OD_chrom,*rpoS −*_*−*OD_chrom,WT_. Arrow labels indicate change in OD_chrom_ of *rpoS*^*−*^ relative to WT. Significance annotations are based on exact two-tailed Wilcoxon signed-rank tests of within-dish differences, with *p* values Bonferroni-adjusted. **, adjusted *p <* 0.01; *, adjusted *p <* 0.05; n.s., not significant; *n* = 11–12 dishes per condition.

All conditions showed measurable CR staining (Fig. 2B). If *σ*^S^ plays a role in biofilm formation, *rpoS*^*−*^ should exhibit altered Congo red binding. The effect we observed, reported as Hodges–Lehmann estimate [95% CI], was medium-dependent: In HS-3 grown on Miller LB, *rpoS* disruption did not significantly affect CR staining (*−*0.024 [*−*0.047, *−*0.011]; *n* = 12, *p*_adj_ = 0.084), whereas on R2A low-nutrient medium, *rpoS* disruption produced a significant drop in CR staining (*−*0.127 [*−*0.139, *−*0.118]; *n* = 12; *p*_adj_ = 0.0020) (Fig. 2B). This is consistent with a *σ*^S^-dependent mechanism for biofilm formation in low-nutrient conditions but not in nutrient-dense environments. In the control group, PA14, *rpoS* disruption was significantly associated with a decrease in CR staining in both Miller LB (*−*0.138 [*−* 0.185, *−*0.092]; *n* = 12; *p*_adj_ = 0.0020) and R2A (*−*0.124 [*−*0.134, *−* 0.110]; *n* = 11; *p*_adj_ = 0.0039), which is expected because *rpoS* is a coordinator of general stress response in PA14 (14) (Fig. 2B).

Given that disruption of *rpoS* significantly reduced CR staining in R2A, our results support a regulatory model in which HS-3’s *σ*^S^ promotes production of biofilm-associated materials under the low-nutrient conditions tested. We did not detect a significant effect of *rpoS* disruption on CR staining under our Miller LB assay conditions.

### *rpoS* disruption does not significantly alter HS-3 survival under tested general environmental stressors

Numerous studies have established the central role of *rpoS* in mediating the survival response to environmental exposures such as heat, hydrogen peroxide (H_2_O_2_), and ethanol (EtOH), in model species such as *Escherichia coli* and *Pseudomonas aeruginosa* (12, 26, 27). If *σ*^S^ plays a role in general stress response, we predicted that the HS-3 *rpoS*^*−*^ mutant should have diminished survival under thermal, oxidative, and envelope stress challenges compared to WT.

After 48 hours of growth at room temperature into stationary phase in R2A, a nutrient-poor medium mimicking the oligotrophic conditions of HS-3’s cave habitat, WT and *rpoS*^*−*^ strains were challenged with either 43.5 °C heat, 0.6 mM H_2_O_2_, or 20% EtOH for around 1 hour (Table 1). Both WT and *rpoS*^*−*^ lost viability steadily throughout each exposure, with heat causing a loss of approximately 0.08 log_10_ CFU mL^*−*1^ min^*−*1^, hydrogen peroxide causing a loss of approximately 0.05 log_10_ CFU mL^*−*1^ min^*−*1^, and EtOH causing a loss of approximately 0.06 log_10_ CFU mL^*−*1^ min^*−*1^ across both WT and *rpoS*^*−*^ strains (Supplementary Fig. 8A–C).

**Table 1.** No significant effect from disruption of *rpoS* on death rate of HS-3 under heat, oxidative, and ethanol stress. Mean *±*SEM of the log-linear death rate fitted to each of *n* = 4 biological replicates per strain. *p*, Welch’s *t*-test on the per-replicate rates. No comparison is significant, before or after Bonferroni correction across all strain pairs.

| Stress | Exposure | Death rate ($\log_{10}$ CFU mL$^{-1}$ min$^{-1}$) | | *rpoS*$^{-}$ vs. WT<br>$\Delta$ (95% CI) | $p$ |
| --- | --- | --- | --- | --- | --- |
| | | WT | *rpoS*$^{-}$ | | |
| Heat | 43.5 °C | $-0.0816 \pm 0.0020$ | $-0.0789 \pm 0.0006$ | $+0.0027 (-0.0034, +0.0089)$ | 0.27 |
| H<sub>2</sub>O<sub>2</sub> | 0.6 mM | $-0.0495 \pm 0.0011$ | $-0.0475 \pm 0.0013$ | $+0.0020 (-0.0023, +0.0063)$ | 0.30 |
| EtOH | 20% v/v | $-0.0583 \pm 0.0056$ | $-0.0587 \pm 0.0027$ | $-0.0004 (-0.0172, +0.0164)$ | 0.95 |

However, there was no significant difference in the death rates between WT and *rpoS*^*−*^ under any of the three stressors. These results contrast with RpoS-dependent resistance to heat and H_2_O_2_ in *P. aeruginosa* (12) and RpoS-dependent resistance to ethanol in *E. coli* (27).

Although RpoS commonly regulates the bacterial general stress response, its physiological role can be more specialized in some bacteria. In *P. aeruginosa*, RpoS regulates the transition between planktonic and biofilm-associated life cycles (14, 15). In *B. burgdorferi* and *V. cholerae*, it regulates host-associated virulence or life cycle transitions (16, 28, 29). These precedents for RpoS as a regulator of life-cycle transitions, along with the apparent narrower scope of *σ*^S^ in HS-3, warrant additional research investigating whether RpoS activity may be associated with phase transition in HS-3’s unique multicellular life cycle.

## Conclusion

A better understanding of the complex life cycle of HS-3, encompassing early multicellular, biofilm-associated growth followed by internal differentiation into motile coccoid cells, may provide insight into how systems arise for the transition from single-to multicellular states in prokaryotic organisms and yield strategies for synthetic multicellular design. However, the genetic tractability of HS-3 was previously unknown, hindering investigation. To assess its tractability, we sought to adapt common cloning techniques to generate the first mutant in HS-3, targeting *rpoS*, which encodes the alternative sigma factor *σ*^S^ commonly associated with regulating general stress response, for insertional disruption.

Here, we present the first mutant strain of HS-3, the *rpoS*^*−*^ strain, and assess its response to general stressors under standardized assay conditions. We demonstrate that, when challenged with nutrient-poor conditions, the *rpoS*^*−*^ strain shows reduced CR binding (*−*0.127 [*−*0.139, *−*0.118]; *n* = 12; *p*_adj_ = 0.0020), suggesting *rpoS* upregulates production of biofilm-associated materials in these conditions.

We also show by disrupting the *rpoS* gene through insertion by homologous recombination that HS-3 is amenable to modification. Although whole-genome sequencing confirmed the insertion, this does not guarantee complete loss of RpoS activity. The insertional inactivation resulted in two defective copies of *rpoS*: one under control of its native promoter but severely truncated, and the other lacking its native promoter, native RBS, and initiator methionine. It is possible, albeit unlikely, that a cryptic promoter and RBS support errant RpoS expression from this second copy. For future work, we aim to establish methods for clean in-frame deletions in HS-3 to avoid such concerns.

Surprisingly, *rpoS* disruption had no detectable effect in response to other general stressors when challenged under traditional assays. Given that HS-3 is a previously uncharacterized environmental isolate, future assays should examine stresses relevant to HS-3’s cave habitat. RpoS may indeed be required for HS-3 to withstand cave-specific stressors yet to be identified. While further tuning of assay conditions may reveal *rpoS*-dependent responses to stresses other than nutrient limitation, there is precedent for *rpoS* regulating a narrower range of stress responses rather than being a master regulator of general stress response (28), as well as several cases of *rpoS* directly controlling major behavioral switches (14, 16). It has been proposed that HS-3’s life cycle involves forming biofilms on cave surfaces in the aftermath of flooding, and our findings motivate the hypothesis that *rpoS* may contribute to biofilm-associated phenotypes under these conditions. Future work should identify the genetic regulators of HS-3’s multicellular life cycle and determine whether RpoS contributes to transitions between stages.

## Methods

### A. Strains and culturing

The unmodified HS-3 strain was provided by Dr. Mizuno and designated parental HS-3(1). An HS-3 *kan*^R^ insertion strain was used as the WT-equivalent control and designated WT, and the HS-3 *rpoS*::*kan*^R^ insertion mutant is designated *rpoS*^*−*^. TransforMax™ EC100D *pir*^+^ Electrocompetent *E. coli* (LGC Biosearch Technologies, Petaluma, CA, USA; #ECP09500) and S17 *λpir* Dap^*−*^

*E. coli* strains were used in cloning (30). *Pseudomonas aeruginosa* PA14 (WT) and *rpoS*::*kan*^R^ (*rpoS*^*−*^) strains, used here as controls, were provided by Dr. Marvin Whiteley (13).

Miller LB and R2A media (0.5 g yeast extract, 0.5 g peptone, 0.5 g casamino acids, 0.5 g glucose, 0.3 g K_2_HPO_4_, 0.05 g MgSO_4_ per liter of DI water) were used. R2A is a low-nutrient medium initially formulated for culturing of oligotrophic bacterial cultures (25). R2A more closely reflects the natural environment of the cave-dwelling HS-3 and is used to induce low-nutrient response in stationary-phase cultures (1).

### B. Generating a disruption mutant in HS-3

The HS-3 *rpoS*^*−*^ strain was constructed using homologous recombination to insert a plasmid encoding a resistance cassette within the coding region of *rpoS*. Operon-mapper was used to confirm that *rpoS* in HS-3 is the final gene within its operon (31). To generate the plasmid for homologous recombination, a two-piece Gibson Assembly was designed using plasmid pRE118 as the backbone (32). A 500 nt internal fragment of *rpoS* was amplified by PCR to exclude the *rpoS* start codon and begin in-frame with the second codon of *rpoS*.

The pRE118 plasmid was chosen because HS-3 does not encode *pir*, which is required for its replication. pRE118 was linearized by XhoI and SphI (NEB) double digestion, and pRE118 and the *rpoS* fragment were amplified by Q5 polymerase (NEB) in 4% DMSO. Gibson assembly was performed using Gibson Assembly® Master Mix (NEB, #E2611L).

This plasmid was electroporated into EC100D *pir*^+^ *E. coli*, isolated using QIAprep® Spin Miniprep Kit (Qiagen, #27106), and electroporated again into S17 *λpir* Dap^*−*^ *E. coli*. The S17 strain was mated with HS-3 on Dap-supplemented LB, and HS-3 integrants were selected on kanamycin lacking Dap.

### C. Sequencing

Strains were inoculated in 50 mL of LB, shaking at 21 °C, and grown for 48 h. Cells from the full culture volume were pelleted at 6,000 *×g* for 5 min. Samples were washed twice in PBS, then suspended in 500 *µ*L of DNA/RNA Shield (Zymo Research, Irvine, CA, USA). Genomic DNA extraction and whole-genome sequencing were performed by Plasmidsaurus using Oxford Nanopore long-read sequencing. Genome assembly and annotation were performed using the Plasmidsaurus analysis pipeline.

### D. Gene identification

To confirm the identity of HS-3 *rpoS* (locus tag AC4KKT_02260) and differentiate it from other alternative sigma factors, including its ancestral *rpoD* (17), each translated sequence was submitted to EBI InterProScan v5.78-109.0 (33), which runs HMMER 3.3 (34), and matched against the Pfam 38.2 database (35) for conserved protein domains. HS-3 alternative sigma factors were compared to homologs from three *γ*-proteobacteria: *Escherichia coli* K-12 (GCF_000005845.2), *Pseudomonas aeruginosa* PA14 (GCF_000014625.1), and *Vibrio cholerae* (GCF_008369605.1).

### E. Minimum inhibitory concentration assay

MICs of kanamycin, chloramphenicol, and gentamicin were characterized in a 96-well plate by inoculating wells with culture that was normalized to OD_600_ = 0.1 and then diluted 200-fold in Miller LB. Wells were then treated with gentamicin, kanamycin, and chloramphenicol to concentrations of 160, 200, and 160 *µ*g/mL, respectively, and to serially two-fold diluted concentrations of each antibiotic. Cells were grown shaking at room temperature for 48 h. MIC was recorded as the lowest concentration that yielded no apparent growth.

### F. Congo red binding assay

Strains were inoculated in Miller LB or R2A and grown shaking at 21 °C. Overnight cultures were normalized to OD_600_ = 0.1. 10 *µ*L aliquots were spotted onto agar plates of the respective medium supplemented with 40 *µ*g/mL Congo red, adapting methods from (21). *P. aeruginosa* strains PA14 WT and *rpoS*^*−*^ were used as controls. Plates were incubated at 21 °C for 72 h before imaging. Images were captured on an iPhone 16 Pro Max in 48 MP RAW format within a light box to minimize spectral variation across images (36).

### G. Congo red quantification

To quantify CR binding, we implemented an application of digital image colorimetry to measure dye-specific chromatic absorption while controlling for non-specific colony opacity and variations in lighting. Because a logarithmic absorbance is valid only on radiometrically linear signal, raw image files (DNG) were decoded into a 16-bit linear-light color space with unity gamma and no brightness adjustments (37).

Colonies were segmented from surrounding agar by color-distance thresholding. To correct for local variations in illumination and exposure, median RGB values were computed in a local annulus of 1.3–1.5 *×* the effective colony radius. Segments and annuli were manually inspected for correct margins.

Following the precedent of converting RGB intensity ratios into apparent optical densities (38, 39), per-channel optical densities were calculated as:

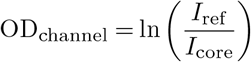

where *I*_core_ is mean intensity within the colony, and *I*_ref_ is median intensity around the annulus, such that a colony with lower intensity within a given channel has greater channel-specific apparent OD (e.g. a colony that strongly reflects red will have a low OD_R_).

We then combined the red and green channels into a chromatic OD:

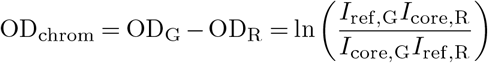

CR absorbs green light and reflects red light, so OD_chrom_ captures both green absorption and red reflection while controlling for the broadband light scattering from increased biomass, such that a greater OD_chrom_ indicates greater CR binding.

Differences were evaluated by exact two-tailed Wilcoxon signed-rank tests on within-dish (mutant - control) differences, and *p* values were adjusted using Bonferroni correction. Effects were reported as Hodges–Lehmann estimates of the location shifts with exact 95% Wilcoxon signed-rank confidence intervals.

### H. Environmental stress response assays

For each assay, strains were inoculated to OD_600_ = 0.1 in 5 mL of R2A medium and grown shaking at 200 rpm at 21 °C for 48 h. To assess resistance to heat, 200 *µ*L samples were incubated in a Mastercycler X50a (Eppendorf, Germany) at 43.5 °C for 80 min. 20 *µ*L aliquots were taken at 0, 20, 40, 60, and 80 min and serially diluted 10-fold in PBS (40). 100 *µ*L of the diluted aliquots were plated on LB agar. To assess resistance to oxidative stress, 1 mL samples were washed in PBS, pelleted at 10,000 *×g* for 3 minutes, re-suspended in 1 mL of 0.6 mM H_2_O_2_ in PBS, and incubated for 80 min. 100 *µ*L aliquots were taken at 0, 20, 40, 60, and 80 min and immediately washed and serially diluted 10-fold in PBS (41), (42). 100 *µ*L aliquots were plated on LB agar. To assess resistance to EtOH, 1 mL samples were washed in PBS, pelleted at 10,000 *×g* for 3 minutes, re-suspended in 1 mL of 20% EtOH in PBS, and incubated for 60 min. 100 *µ*L aliquots were taken at 0, 10, 20, 40, and 60 min and serially diluted 10-fold in PBS. 100 *µ*L aliquots were plated on LB agar (43). After each assay, the LB plates were incubated at 21 °C for 48 h, and then CFU counts were recorded. For each assay, WT and *rpoS*^*−*^ strains were each represented by 4 biological replicates. A log-linear death rate was fitted by least squares to each biological replicate, and strains were compared by Welch’s *t*-test.

## Data Availability Statement

The data underlying this article are available in the article and in its online supplementary material. Code used in this study is available at https://github.com/Zbedd/HS-3_rpoS_analysis.

Genome assemblies have been deposited in GenBank under the BioProject PRJNA1480197 for HS-3 wild type (BioSample SAMN61019079, accession JCEHQB000000000) and *rpoS*^*−*^ (BioSample SAMN61019081, accession JCE-HQA000000000) strains.

## Contributions

Z.B.: Conceptualization, Methodology, Investigation, Formal analysis, Software, Data curation, Visualization, Writing – original draft, Writing – review & editing. I.C.: Conceptualization, Methodology, Investigation, Formal analysis, Software, Visualization, Writing – original draft, Writing – review & editing. C.Z.: Conceptualization, Methodology, Writing – review & editing, Supervision. B.H.: Conceptualization, Methodology, Supervision, Resources, Funding acquisition.

## Acknowledgments

We thank Dr. Kouhei Mizuno for providing the HS-3 strain used in this work, the Whiteley Lab for providing the control strains used in this work, Parker Smith for providing the apparatus used for Congo red imaging, and Dr. William Ratcliff for providing for helpful discussions and valuable insights. We acknowledge support from the Georgia Institute of Technology President’s Undergraduate Research Awards (PURA) to I.C. and Z.B., the Georgia Tech School of Biological Sciences to B.K.H., and a U.S. Department of Education GAANN Fellowship (award P200A210046) to C.Z.

AI was used to assist with the development and troubleshooting of analysis code and manuscript polishing. The authors reviewed all AI-assisted work and take responsibility for the final manuscript.

## Notes

### Competing Interest Statement

The authors have declared no competing interest.

## Bibliography

1. Kouhei Mizuno, Mais Maree, Toshihiko Nagamura, Akihiro Koga, Satoru Hirayama, Soichi Furukawa, Kenji Tanaka, and Kazuya Morikawa. Novel multicellular prokaryote discovered next to an underground stream. eLife, 11:e71920, October 2022. ISSN 2050-084X. doi: 10.7554/eLife.71920.

2. Daniel C. Volke, Enrico Orsi, and Pablo I. Nikel. Emergent CRISPR–Cas-based technologies for engineering non-model bacteria. Current Opinion in Microbiology, 75:102353, October 2023. ISSN 1369-5274. doi: 10.1016/j.mib.2023.102353.

3. Amanda T. Alker, Morgan V. Farrell, Alpher E. Aspiras, Tiffany L. Dunbar, Andriy Fedoriouk, Jeffrey E. Jones, Sama R. Mikhail, Gabriella Y. Salcedo, Bradley S. Moore, and Nicholas J. Shikuma. A modular plasmid toolkit applied in marine bacteria reveals functional insights during bacteria-stimulated metamorphosis. mBio, 14(4):e0150223, August 2023. ISSN 2150-7511. doi: 10.1128/mbio.01502-23.

4. Zachary Jansen, Abdulaziz Alameri, Qiyao Wei, Devon L. Kulhanek, Andrew R. Gilmour, Sean Halper, Nathan D. Schwalm, and Ross Thyer. A modular toolkit for environmental Rhodococcus, Gordonia, and Nocardia enables complex metabolic manipulation. Applied and Environmental Microbiology, 90(8):e00340–24, July 2024. doi: 10.1128/aem.00340-24.

5. Christopher D. Johnston, Sean L. Cotton, Susan R. Rittling, Jacqueline R. Starr, Gary G. Borisy, Floyd E. Dewhirst, and Katherine P. Lemon. Systematic evasion of the restriction-modification barrier in bacteria. Proceedings of the National Academy of Sciences, 116(23): 11454–11459, June 2019. doi: 10.1073/pnas.1820256116.

6. Leen De Gelder, JoséM Ponciano, Paul Joyce, and Eva M Top. Stability of a promiscuous plasmid in different hosts: no guarantee for a long-term relationship. Microbiology, 153(2): 452–463, 2007. ISSN 1465-2080. doi: 10.1099/mic.0.2006/001784-0.

7. Ewa Lewicka, Patrycja Dolowy, Jolanta Godziszewska, Emilia Litwin, Marta Ludwiczak, and Grazyna Jagura-Burdzy. Transcriptional Organization of the Stability Module of Broad-Host-Range Plasmid RA3, from the IncU Group. Applied and Environmental Microbiology, 86 (16):e00847–20, August 2020. doi: 10.1128/AEM.00847-20.

8. Marita Torrissen Mårli, Oddvar Oppegaard, Davide Porcellato, Daniel Straume, and Morten Kjos. Genetic modification of Streptococcus dysgalactiae by natural transformation. mSphere, 9(7):e00214–24, June 2024. doi: 10.1128/msphere.00214-24.

9. Regine Hengge-Aronis. Signal Transduction and Regulatory Mechanisms Involved in Control of the σS (RpoS) Subunit of RNA Polymerase. Microbiology and Molecular Biology Reviews, 66(3):373–395, September 2002. ISSN 1092-2172. doi: 10.1128/MMBR.66.3.373-395.2002.

10. Sarah M. Chiang and Herb E. Schellhorn. Evolution of the RpoS regulon: origin of RpoS and the conservation of RpoS-dependent regulation in bacteria. Journal of Molecular Evolution, 70(6):557–571, June 2010. ISSN 1432-1432. doi: 10.1007/s00239-010-9352-0.

11. Aurelia Battesti, Nadim Majdalani, and Susan Gottesman. The RpoS-mediated general stress response in Escherichia coli. Annual Review of Microbiology, 65:189–213, 2011. ISSN 1545-3251. doi: 10.1146/annurev-micro-090110-102946.

12. S. J. Suh, L. Silo-Suh, D. E. Woods, D. J. Hassett, S. E. West, and D. E. Ohman. Effect of rpoS mutation on the stress response and expression of virulence factors in Pseudomonas aeruginosa. Journal of Bacteriology, 181(13):3890–3897, July 1999. ISSN 0021-9193. doi: 10.1128/JB.181.13.3890-3897.1999.

13. M. Whiteley, M. R. Parsek, and E. P. Greenberg. Regulation of quorum sensing by RpoS in Pseudomonas aeruginosa. Journal of Bacteriology, 182(15):4356–4360, August 2000. ISSN 0021-9193. doi: 10.1128/JB.182.15.4356-4360.2000.

14. Sophie Bouillet, Moly Ba, Laetitia Houot, Chantal Iobbi-Nivol, and Christophe Bordi. Connected partner-switches control the life style of Pseudomonas aeruginosa through RpoS regulation. Scientific Reports, 9(1):6496, April 2019. ISSN 2045-2322. doi: 10.1038/s41598-019-42653-5.

15. Clayton W. Hall, Aaron J. Hinz, Luke B.-P. Gagnon, Li Zhang, Jean-Paul Nadeau, Sarah Copeland, Bratati Saha, and Thien-Fah Mah. Pseudomonas aeruginosa Biofilm Antibiotic Resistance Gene ndvB Expression Requires the RpoS Stationary-Phase Sigma Factor. Applied and Environmental Microbiology, 84(7):e02762–17, April 2018. ISSN 1098-5336. doi: 10.1128/AEM.02762-17.

16. Alex Toftgaard Nielsen, Nadia A. Dolganov, Glen Otto, Michael C. Miller, Cheng Yen Wu, and Gary K. Schoolnik. RpoS Controls the Vibrio cholerae Mucosal Escape Response. PLOS Pathogens, 2(10):e109, October 2006. ISSN 1553-7374. doi: 10.1371/journal.ppat.0020109.

17. Herb E. Schellhorn. Function, Evolution, and Composition of the RpoS Regulon in Escherichia coli. Frontiers in Microbiology, 11, September 2020. ISSN 1664-302X. doi: 10.3389/fmicb.2020.560099.

18. Xiaoxiang Liu, Jun Xu, Junli Zhu, Peng Du, and Aihua Sun. Combined Transcriptome and Proteome Analysis of RpoS Regulon Reveals Its Role in Spoilage Potential of Pseudomonas fluorescens. Frontiers in Microbiology, 10, February 2019. ISSN 1664-302X. doi: 10.3389/fmicb.2019.00094.

19. Yasuhiko Irie, Melissa Starkey, Adrianne N Edwards, Daniel J Wozniak, Tony Romeo, and Matthew R Parsek. Pseudomonas aeruginosa biofilm matrix polysaccharide Psl is regulated transcriptionally by RpoS and post-transcriptionally by RsmA. Molecular Microbiology, 78 (1):158–172, October 2010. ISSN 0950-382X. doi: 10.1111/j.1365-2958.2010.07320.x.

20. Kelly M. Colvin, Vernita D. Gordon, Keiji Murakami, Bradley R. Borlee, Daniel J. Wozniak, Gerard C. L. Wong, and Matthew R. Parsek. The Pel Polysaccharide Can Serve a Structural and Protective Role in the Biofilm Matrix of Pseudomonas aeruginosa. PLOS Pathogens, 7 (1):e1001264, January 2011. ISSN 1553-7374. doi: 10.1371/journal.ppat.1001264.

21. Karishma Bisht, Alex R. Luecke, and Catherine A. Wakeman. Temperature-specific adaptations and genetic requirements in a biofilm formed by Pseudomonas aeruginosa. Frontiers in Microbiology, 13, January 2023. ISSN 1664-302X. doi: 10.3389/fmicb.2022.1032520.

22. Jens Bo Andersen, Louise Dahl Hultqvist, Charlotte Uldahl Jansen, Tim Holm Jakobsen, Martin Nilsson, Morten Rybtke, Jesper Uhd, Blaine Gabriel Fritz, Roland Seifert, Jens Berthelsen, Thomas Eiland Nielsen, Katrine Qvortrup, Michael Givskov, and Tim Tolker-Nielsen. Identification of small molecules that interfere with c-di-GMP signaling and induce dispersal of Pseudomonas aeruginosa biofilms. npj Biofilms and Microbiomes, 7(1):59, July 2021. ISSN 2055-5008. doi: 10.1038/s41522-021-00225-4.

23. Marianyoly Ortiz, Antje Legatzki, Julia W Neilson, Brandon Fryslie, William M Nelson, Rod A Wing, Carol A Soderlund, Barry M Pryor, and Raina M Maier. Making a living while starving in the dark: metagenomic insights into the energy dynamics of a carbonate cave. The ISME Journal, 8(2):478–491, February 2014. ISSN 1751-7362. doi: 10.1038/ismej.2013.159.

24. Suprokash Koner, Jung-Sheng Chen, Bing-Mu Hsu, Chao-Wen Tan, Cheng-Wei Fan, Tsung-Hsien Chen, Bashir Hussain, and Viji Nagarajan. Assessment of Carbon Substrate Catabolism Pattern and Functional Metabolic Pathway for Microbiota of Limestone Caves. Microorganisms, 9(8):1789, August 2021. ISSN 2076-2607. doi: 10.3390/microorganisms9081789.

25. D J Reasoner and E E Geldreich. A new medium for the enumeration and subculture of bacteria from potable water. Applied and Environmental Microbiology, 49(1):1–7, January 1985. ISSN 0099-2240. doi: 10.1128/aem.49.1.1-7.1985.

26. Andrew J. Hryckowian and Rodney A. Welch. RpoS Contributes to Phagocyte Oxidase-Mediated Stress Resistance during Urinary Tract Infection by Escherichia coli CFT073. mBio, 4(1):https://doi.org/10.1128/mbio.00023–13, February 2013. doi: 10.1128/mbio.00023-13.

27. Anne Farewell, Kristian Kvint, and Thomas Nyström. uspB, a New ςS-Regulated Gene in Escherichia coli Which Is Required for Stationary-Phase Resistance to Ethanol. Journal of Bacteriology, 180(23):6140–6147, December 1998. doi: 10.1128/jb.180.23.6140-6147.1998.

28. Melissa J. Caimano, Christian H. Eggers, Karsten R. O. Hazlett, and Justin D. Radolf. RpoS is not central to the general stress response in Borrelia burgdorferi but does control expression of one or more essential virulence determinants. Infection and Immunity, 72(11): 6433–6445, November 2004. ISSN 0019-9567. doi: 10.1128/IAI.72.11.6433-6445.2004.

29. A. Hübner, X. Yang, D. M. Nolen, T. G. Popova, F. C. Cabello, and M. V. Norgard. Expression of Borrelia burgdorferi OspC and DbpA is controlled by a RpoN-RpoS regulatory pathway. Proceedings of the National Academy of Sciences of the United States of America, 98(22): 12724–12729, October 2001. ISSN 0027-8424. doi: 10.1073/pnas.231442498.

30. R. Simon, U. Priefer, and A. Pühler. A Broad Host Range Mobilization System for In Vivo Genetic Engineering: Transposon Mutagenesis in Gram Negative Bacteria. Bio/Technology, 1(9):784–791, November 1983. ISSN 1546-1696. doi: 10.1038/nbt1183-784.

31. Blanca Taboada, Karel Estrada, Ricardo Ciria, and Enrique Merino. Operon-mapper: a web server for precise operon identification in bacterial and archaeal genomes. Bioinformatics, 34(23):4118–4120, December 2018. ISSN 1367-4803. doi: 10.1093/bioinformatics/bty496.

32. R. A. Edwards, L. H. Keller, and D. M. Schifferli. Improved allelic exchange vectors and their use to analyze 987P fimbria gene expression. Gene, 207(2):149–157, January 1998. ISSN 0378-1119. doi: 10.1016/s0378-1119(97)00619-7.

33. Philip Jones, David Binns, Hsin-Yu Chang, Matthew Fraser, Weizhong Li, Craig McAnulla, Hamish McWilliam, John Maslen, Alex Mitchell, Gift Nuka, Sebastien Pesseat, Antony F. Quinn, Amaia Sangrador-Vegas, Maxim Scheremetjew, Siew-Yit Yong, Rodrigo Lopez, and Sarah Hunter. InterProScan 5: genome-scale protein function classification. Bioinformatics, 30(9):1236–1240, May 2014. ISSN 1367-4811. doi: 10.1093/bioinformatics/btu031.

34. Sean R. Eddy. Accelerated Profile HMM Searches. PLOS Computational Biology, 7(10):e1002195, October 2011. ISSN 1553-7358. doi: 10.1371/journal.pcbi.1002195.

35. Typhaine Paysan-Lafosse, Antonina Andreeva, Matthias Blum, Sara Rocio Chuguransky, Tiago Grego, Beatriz Lazaro Pinto, Gustavo A Salazar, Maxwell L Bileschi, Felipe LlinaresLópez, Laetitia Meng-Papaxanthos, Lucy J Colwell, Nick V Grishin, R Dustin Schaeffer, Damiano Clementel, Silvio C E Tosatto, Erik Sonnhammer, Valerie Wood, and Alex Bateman. The Pfam protein families database: embracing AI/ML. Nucleic Acids Research, 53 (D1):D523–D534, November 2024. ISSN 0305-1048. doi: 10.1093/nar/gkae997.

36. Parker Smith and Martin Schuster. Inexpensive Apparatus for High-Quality Imaging of Microbial Growth on Agar Plates. Frontiers in Microbiology, 12, June 2021. ISSN 1664-302X. doi: 10.3389/fmicb.2021.689476.

37. Olivier Burggraaff, Norbert Schmidt, Jaime Zamorano, Klaas Pauly, Sergio Pascual, Carlos Tapia, Evangelos Spyrakos, and Frans Snik. Standardized spectral and radiometric calibration of consumer cameras. Optics Express, 27(14):19075–19101, July 2019. ISSN 1094-4087. doi: 10.1364/OE.27.019075.

38. A. C. Ruifrok and D. A. Johnston. Quantification of histochemical staining by color deconvolution. Analytical and Quantitative Cytology and Histology, 23(4):291–299, August 2001.

39. Yoshiki Soda, Kye J. Robinson, Thomas J. Cherubini, and Eric Bakker. Colorimetric absorbance mapping and quantitation on paper-based analytical devices. Lab on a Chip, 20 (8):1441–1448, April 2020. ISSN 1473-0197. doi: 10.1039/d0lc00028k.

40. Lucinda Notley-McRobb, Thea King, and Thomas Ferenci. rpoS Mutations and Loss of General Stress Resistance in Escherichia coli Populations as a Consequence of Conflict between Competing Stress Responses. Journal of Bacteriology, 184(3):806–811, February 2002. ISSN 0021-9193. doi: 10.1128/JB.184.3.806-811.2002.

41. Yifan Xu, Andreas Itzek, and Jens Kreth. Comparison of genes required for H2O2 resistance in Streptococcus gordonii and Streptococcus sanguinis. Microbiology, 160(Pt 12):2627– 2638, December 2014. ISSN 1350-0872. doi: 10.1099/mic.0.082156-0.

42. Alexandro Rodríguez-Rojas, Joshua Jay Kim, Paul R. Johnston, Olga Makarova, Murat Eravci, Christoph Weise, Regine Hengge, and Jens Rolff. Non-lethal exposure to H2O2 boosts bacterial survival and evolvability against oxidative stress. PLoS Genetics, 16(3):e1008649, March 2020. ISSN 1553-7390. doi: 10.1371/journal.pgen.1008649.

43. Frieda Jørgensen, Marc Bally, Virginie Chapon-Herve, Gerard Michel, Andrée Lazdunski, Paul Williams, and G. S. A. B. Stewart. RpoS-dependent stress tolerance in Pseudomonas aeruginosa. Microbiology, 145(4):835–844, 1999. ISSN 1465-2080. doi: 10.1099/13500872-145-4-835.

