## Supplemental Data for "Establishing genetic tractability in *Jeongeupia sacculi* HS-3: disruption of *rpoS* hinders biofilm formation"

### Supplementary Information

#### Supplementary Tables

**Table 1. Minimum inhibitory concentrations (MIC, $\mu$g/mL) of HS-3 (parental) and *E. coli* (S17 $\lambda$*pir* Dap$^{-}$), determined by twofold dilution.** MIC is the lowest concentration fully inhibiting growth. Values were consistent across replicates. n.t., not tested.

| Antibiotic | HS-3 | *E. coli* | *n* |
|---|---|---|---|
| Gentamicin | 160 | n.t. | 3 |
| Kanamycin | 100 | 25 | 4 |
| Chloramphenicol | 10 | 20 | 4 |

DRAFT

**Table 2. Oligonucleotides used in this study.** Lowercase denotes pRE118-derived sequence; uppercase denotes HS-3 *rpoS*-derived sequence.

| Sequence (5′–3′) | Length | Role |
|---|---|---|
| aatgccgatatcctattggcAACGATCAAATCGATATGCTGG | 42 nt | *rpoS* F |
| cttttatatagttcatatggTTGATGACATGCACCGG | 37 nt | *rpoS* R |
| TTGCCGGTGCATGTCATCAAccatatgaactatataaaagcaggc | 45 nt | pRE118 F |
| AGCATATCGATTTGATCGTTgccaataggatatcggcatt | 40 nt | pRE118 R |

**Table 3. Bacterial strains used in this study.**

| Strain/source | Background | Relevant genotype | Label |
|---|---|---|---|
| ZB001 | *J. sacculi* HS-3 | Wild type | parental |
| ZB017 | ZB001 | *rpoS*::$\Omega$ *kan*$^{R}$ | *rpoS*$^{-}$ |
| ZB018 | ZB001 | *kan*$^{R}$, random insertion | WT |
| Whiteley et al., 2000 | *P. aeruginosa* | Wild type | WT |
| Whiteley et al., 2000 | *P. aeruginosa* | *rpoS*::$\Omega$ *kan*$^{R}$ | *rpoS*$^{-}$ |

**Table 4. Plasmids used in this study.**

| Plasmid | Source | Relevant characteristics |
|---|---|---|
| pRE118 | Edwards et al., 1998 | Allelic exchange suicide vector; R6K$\gamma$ *ori* ($\lambda$*pir*-dependent), *oriT*, *sacB*, *kan*$^{R}$ |
| pRE118-*rpoS*Flanks | This study | pRE118 carrying a 500 nt internal fragment of HS-3 *rpoS* |

#### Supplementary Figures

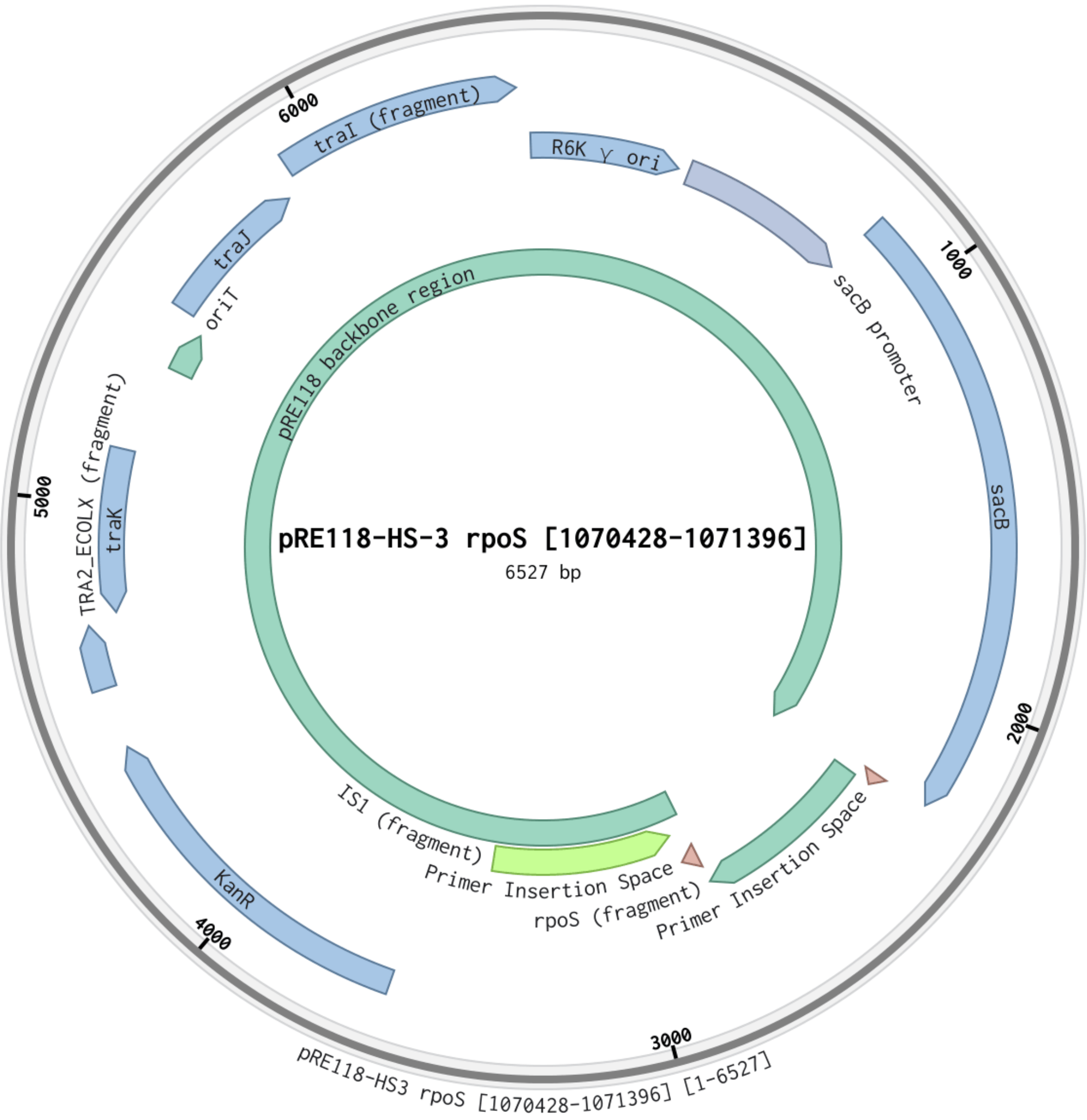


**Fig. 1. Plasmid map of pRE118-HS-3 *rpoS* [1070428–1071396] used to produce the *rpoS*$^{-}$ mutant.** Plasmid map produced in Benchling software.

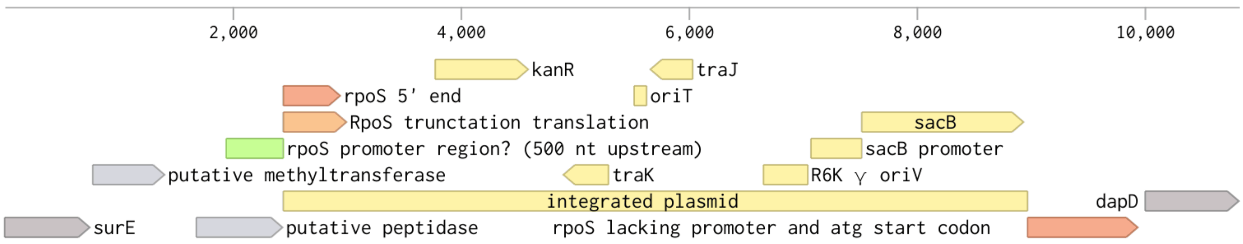


**Fig. 2. Map of pRE118-HS-3 *rpoS* [1070428–1071396] integration into HS-3.** Plasmid map produced in Benchling software.

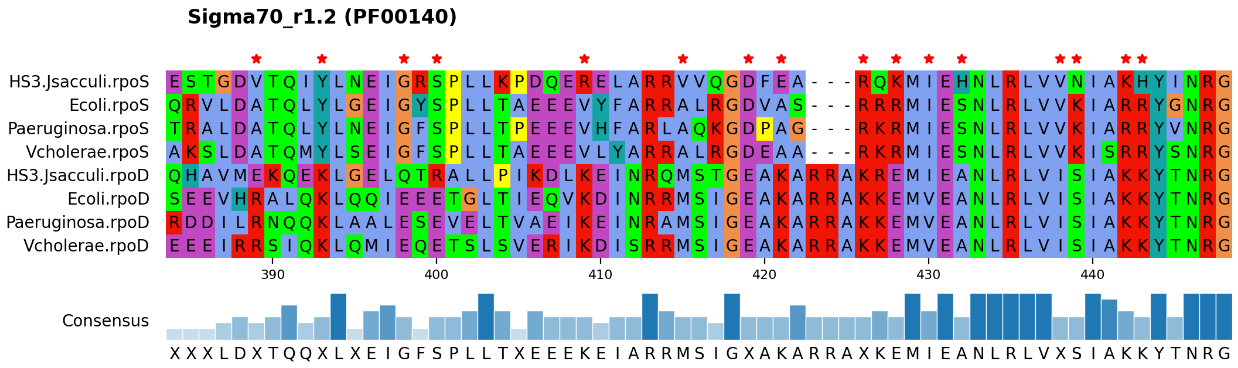


**Fig. 3. Multiple sequence alignment of $\sigma^{70}$ region 1.2 (Pfam PF00140) for RpoS and RpoD across HS-3 and four reference organisms, E. coli, *P. aeruginosa*, *V. cholerae*, and *C. violaceum*.** Red asterisks mark columns where *rpoS* and *rpoD* reference groups converge on different residues, each with $\geq 60\%$ within-group agreement. Residues are colored by Clustal color scheme. The bottom row depicts position-wise consensus.

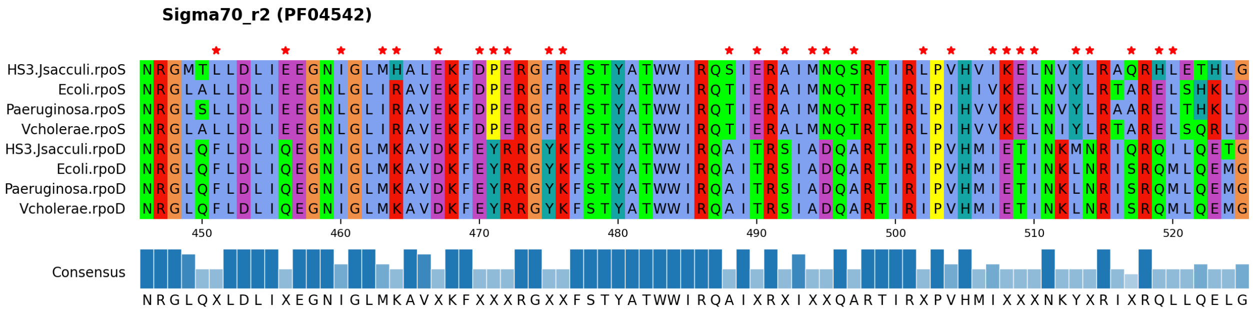


**Fig. 4. Multiple sequence alignment of $\sigma^{70}$ region 2 (Pfam PF04542) for RpoS and RpoD across HS-3 and four reference organisms, E. coli, *P. aeruginosa*, *V. cholerae*, and *C. violaceum*.** Red asterisks mark columns where *rpoS* and *rpoD* reference groups converge on different residues, each with $\geq 60\%$ within-group agreement. Residues are colored by Clustal color scheme. The bottom row depicts position-wise consensus.

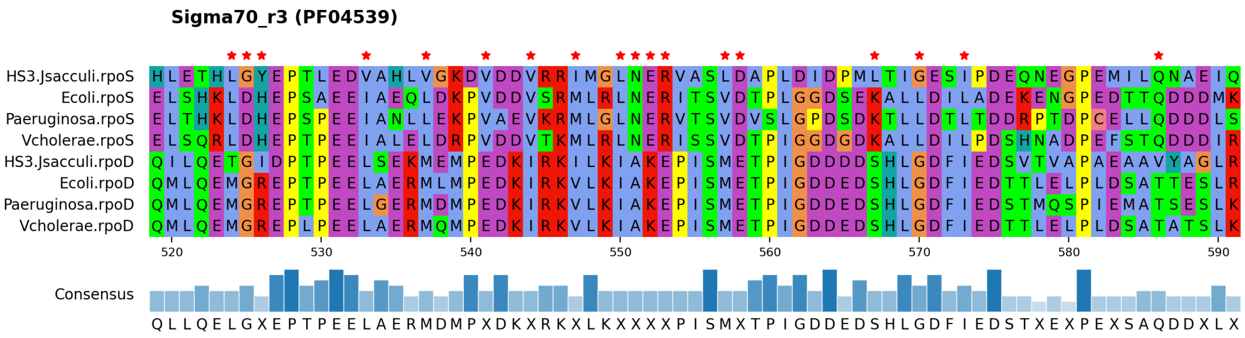


**Fig. 5. Multiple sequence alignment of $\sigma^{70}$ region 3 (Pfam PF04539) for RpoS and RpoD across HS-3 and four reference organisms, E. coli, *P. aeruginosa*, *V. cholerae*, and *C. violaceum*.** Red asterisks mark columns where *rpoS* and *rpoD* reference groups converge on different residues, each with $\geq 60\%$ within-group agreement. Residues are colored by Clustal color scheme. The bottom row depicts position-wise consensus.

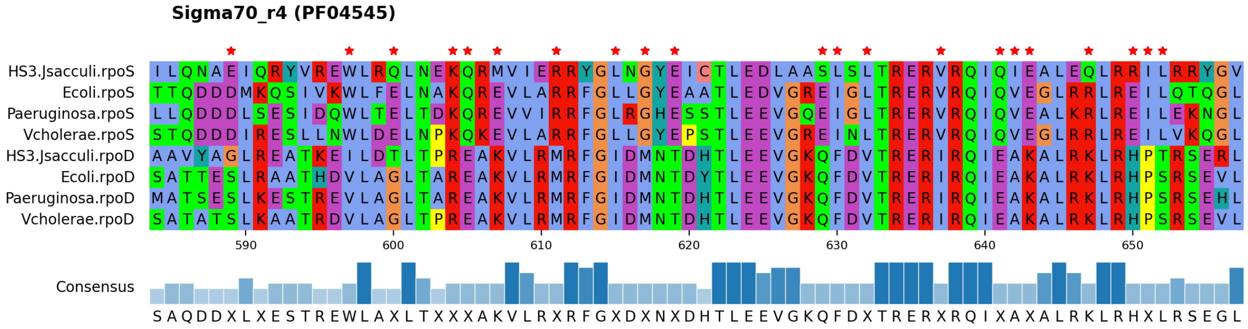

**Fig. 6. Multiple sequence alignment of $\sigma^{70}$ region 4 (Pfam PF04545) for RpoS and RpoD across HS-3 and four reference organisms, E. coli, *P. aeruginosa*, *V. cholerae*, and *C. violaceum*.** Red asterisks mark columns where *rpoS* and *rpoD* reference groups converge on different residues, each with $\geq 60\%$ within-group agreement. Residues are colored by Clustal color scheme. The bottom row depicts position-wise consensus.

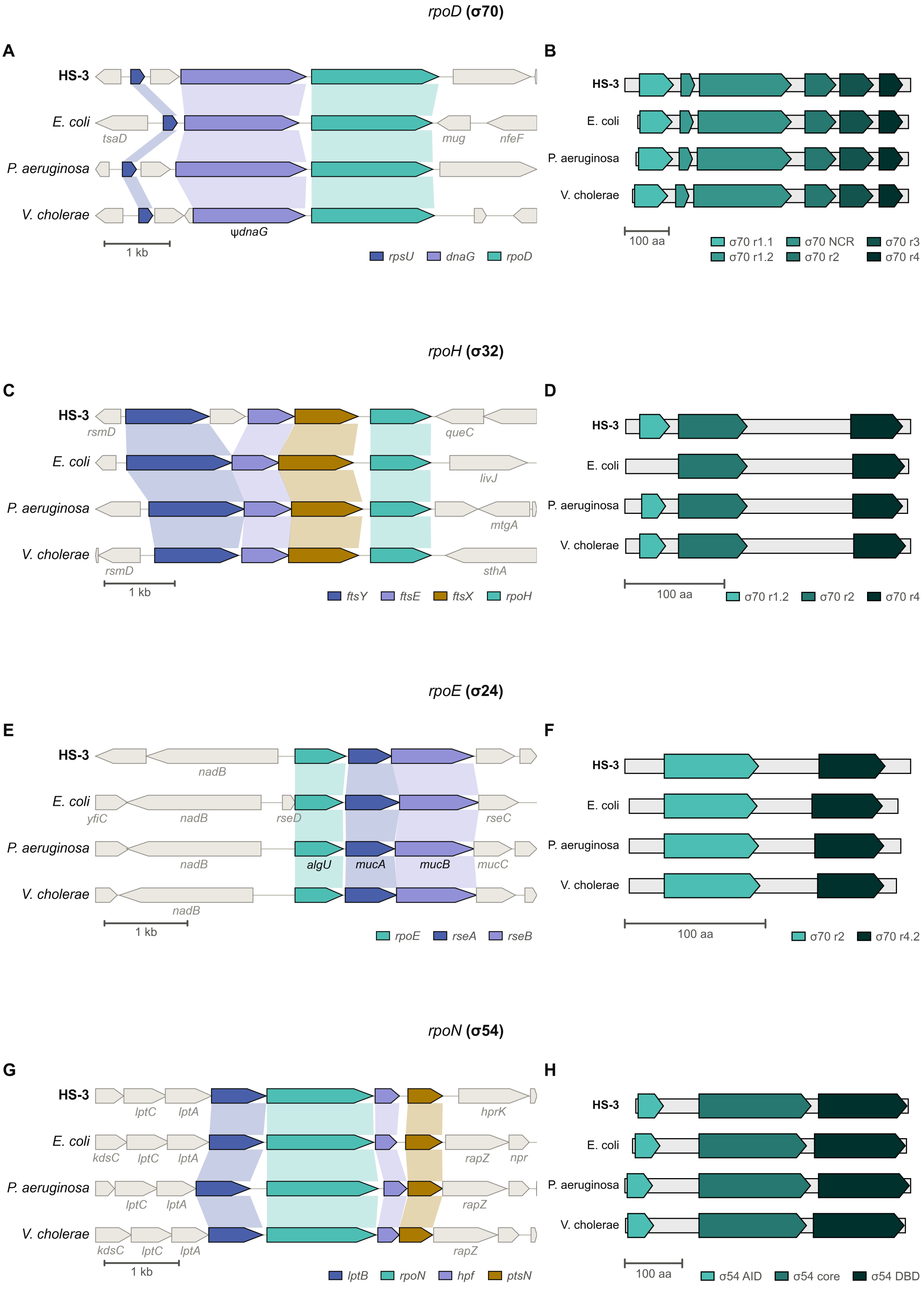


**Fig. 7. Genomic neighborhoods and protein domain architectures for *rpoD* ($\sigma^{70}$) (A, B), *rpoH* ($\sigma^{32}$) (C, D), *rpoE* ($\sigma^{24}$) (E, F), and *rpoN* ($\sigma^{54}$) (G, H) across HS-3 and four reference organisms, E. coli, *P. aeruginosa*, *V. cholerae*, and *C. violaceum*.** Sequences were submitted to EBI InterProScan v5.78-109.0 and matched against the Pfam 38.2 database.

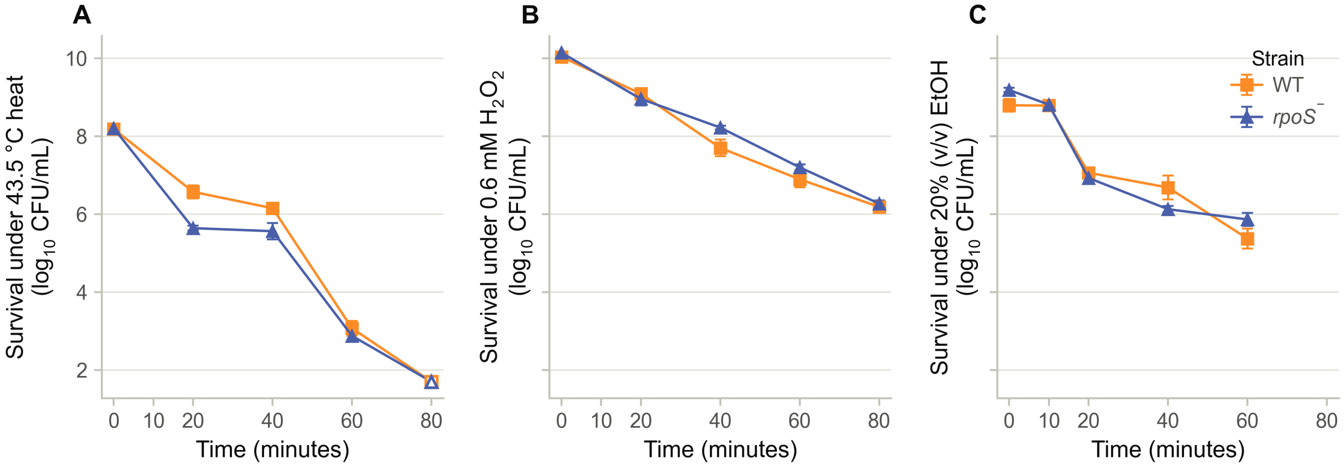


**Fig. 8. Effect of *rpoS* disruption on survival in response to $\mathrm{H_2O_2}$ exposure (0.6 mM).** Time-kill curves for HS-3 WT (orange squares) and *rpoS*$^-$ (blue triangles) exposed to (A) heat, 43.5 °C; (B) $\mathrm{H_2O_2}$, 0.6 mM; and (C) EtOH, 20% v/v. Counts are plotted as $\log_{10}(\mathrm{CFU/mL})$. Points are the mean of $n = 4$ biological replicates. Error bars show ±1 standard error of the mean (SEM), computed on $\log_{10}$-transformed counts. Open markers indicate a timepoint where at least one replicate had no detected CFUs. Strains were compared by Welch's $t$-test of death-rate slopes, and no differences between strains were statistically significant.
